# Temporal regulation of G2 phase avoids therapy-induced senescence caused by DNA replication stress-inducing drugs and provides synergistic cytotoxicity

**DOI:** 10.64898/2026.05.06.723184

**Authors:** Kentaro Nonaka, Takeshi Wakasa, Hiroaki Ochiiwa, Yuki Kataoka, Koji Ando, Eiji Oki, Tomoharu Yoshizumi, Yoshihiko Maehara, Hiroyuki Kitao, Makoto Iimori

**Author notes:** Corresponding authors: Makoto Iimori and Hiroyuki Kitao Oral Medicine Research Center, Fukuoka Dental College, 2-15-1 Tamura, Sawara-ku, Fukuoka, 814-0193, Japan., (ext. 4195 (MI) or 1683 (HK)), (MI) (HK).

## Abstract

The cellular response to DNA replication stress (DRS) provoked by anticancer drugs involves activation of the G2/M checkpoint (which promotes transient cell cycle arrest at G2 phase) and DNA repair, followed by induction of apoptosis or senescence. Here, we activated the p53-p21 pathway and ATR using DRS-inducing drugs, and found that that the transition to senescence depends on the duration of the G2 phase. Shortening of G2 duration by G2/M checkpoint inhibitors led not only to a switch in cell fate from senescence to mitotic entry, but also to effective cell death through carry-over of chromosomal aberrations (generated by DRS-inducing drugs) into mitosis and subsequent mitotic progression. Such enhanced cell death was also observed in p53 deficient cells, which do not normally undergo senescence. Thus, we propose that temporal regulation of G2 phase is an approach to enhancing the effects of DRS-inducing drugs in a manner that is independent of p53 status.

## INTRODUCTION

The cellular response to DNA replication stress (DRS)-inducing drugs involves activation of the G2/M checkpoint, which promotes transient cell-cycle arrest at G2 phase as well as DNA repair, followed by induction of apoptosis or senescence ^1^. Key factors for induction and maintenance of G2 phase arrest in response to DNA damage by DRS-inducing drugs include G2/M checkpoint proteins such as atexia telangiectasia and Rad3-related (ATR) kinase, checkpoint kinase 1 (CHK1) kinase and WEE1 kinase, and the tumor suppressor protein p53. Although ATR plays a crucial role in responses to various types of genotoxic stress, it is activated via generation of extensive single-stranded DNA (ssDNA) breaks, which are coated by replication protein A (RPA) at stalled DNA replication forks, or resected double-stranded DNA ends ^2^. During G2 phase, phospho-activation of CHK1 by ATR inhibits CDC25C phosphatase and increases activity of WEE1 kinase, which negatively regulates cyclin-dependent kinase 1 (CDK1), leading to activation of the G2/M checkpoint^3^.

In another cascade that is responsible for G2/M checkpoint activation through negative regulation of CDK1, p53 contributes to inhibition of CDK1 or nuclear exclusion of CDC25C via downstream factors such as p21 or 14-3-3, respectively^4,5^. By contrast, p53 proficient cells are often induced to undergo cellular senescence by treatment with chemotherapeutic drugs such as DRS-inducing drugs, a process called therapy-induced senescence (TIS) ^6^. The mechanism underlying induction of TIS involves activation of the p53-p21 pathway during G2 phase, leading to nuclear retention and subsequent degradation of cyclin B1 and premature activation of APC/C^Cdh1^ (anaphase-promoting complex/cyclosome and its coactivator Cdh1), as well as pRb-dependent transcriptional repression of mitotic regulators. Such cells exhibit a mitosis skip and exit the cell cycle permanently ^7,8^. Therefore, p53 plays a role both in activating the G2/M checkpoint and in inducing senescence during G2 phase; however, it is unclear how cells in which p53 is activated by DRS during G2 phase are selected to maintain G2 arrest or transition to senescence. Furthermore, we do not know how G2/M checkpoint factors such as ATR, CHK1, and WEE1 are involved in decisions regarding these cell fates.

Although different types of DRS-inducing drugs cause TIS, the relationship between temporal regulation of G2 phase and resulting cell fate, and the effect of p53 status on this relationship, has not been explored. Here, we reveal how cell fate induced by TIS via mitosis skip is determined during G2 phase, and show that avoiding TIS caused by DRS-inducing drugs can lead to induction of synergistic cytotoxicity.

## RESULTS

### ATR activity is required for prolonged G2 phase duration-dependent transition to senescence

Our previous studies revealed that a DRS-inducing drug, a fluorinated thymidine analogue trifluridine (hereafter referred to as FTD), induced formation of γH2AX and RPA32 nuclear foci in HCT116 cells, indicating the presence of DNA double-strand breaks (DSBs) and ssDNA accumulation; this was accompanied by prolongation of the S-G2 phase ^9,10^. Therefore, to assess such DNA damage responses during S and G2 phase in detail, we synchronized HCT116 cells by inducing cell cycle arrest using thymidine and RO-3306 (a CDK1 inhibitor). We then added FTD to the cell culture medium 5 h after release from RO-3306, when most cells were in G1 or early S phase (Fig. 1A and supplementary Fig.3A) and noted that FTD induced accumulation of ssDNA and activated ATR, as indicated by phosphorylation of RPA32 Ser33 and autophosphorylation of ATR Thr1989 ^11,12^, respectively; this was sustained not only during S phase but also during G2 phase (Fig. 1B). To elucidate the biological significance of ATR activation in FTD-treated cells during G2 phase, we inhibited ATR kinase activity or depleted ATR using the ATR inhibitor (hereafter referred to as ATRi) ceralasertib (AZD6738) or siRNA, respectively. In addition, we added the ATRi to cells 25 h after FTD treatment (a time when numerous cells were just about to enter G2 phase) to restrict the effect of ATR inhibition during G2 (data not shown). Both the ATRi and siRNA increased the number of multinuclear cells rather than decreasing the number of mononuclear cell (Fig. 1C, D, and Supplementary Fig. 1A, B). Previous studies report that FTD induces formation of multinuclear p53-knockout (hereafter referred to as p53-KO) and p53-gain-of-function missense mutant cells, but not that of p53-wild-type cells due to transition into senescence ^9,10^. Consequently, our finding that suppressing ATR activity during FTD treatment induces formation of multinuclear p53-wild-type cells led us to speculate that ATR-inhibits escape of these cells from senescence transition and allows them to enter mitosis. To test this hypothesis, we examined the transition from senescence to mitotic entry and found that addition of ATRi altered the fate of FTD-treated Fucci-SA^13^ cells from mitosis skip, which is a marker of senescence transition characterized by entry into the next G1 phase without separation into two daughter cells^8,9^, to mitotic entry (Fig. 1E, F). Because TIS transition is decided during G2 phase ^7,8^, we performed live-cell imaging to measure the duration of G2 phase in FTD-treated HCT116 cells harboring PIP-Fucci, and found that the G2 phase was much longer in cells that skip mitosis than in those that enter to mitosis. Notably, addition of the ATRi to FTD shortened the duration of G2 phase significantly; indeed, it was then the same as that in unchallenged cells (Fig. 1G, H). Taken together, these data indicate that activation of ATR during in FTD-treated cells during G2 phase is necessary for transition to senescence via a prolonged G2 phase, and that shortening G2 switches cell fate from senescence to mitotic entry.

**Figure 1.**
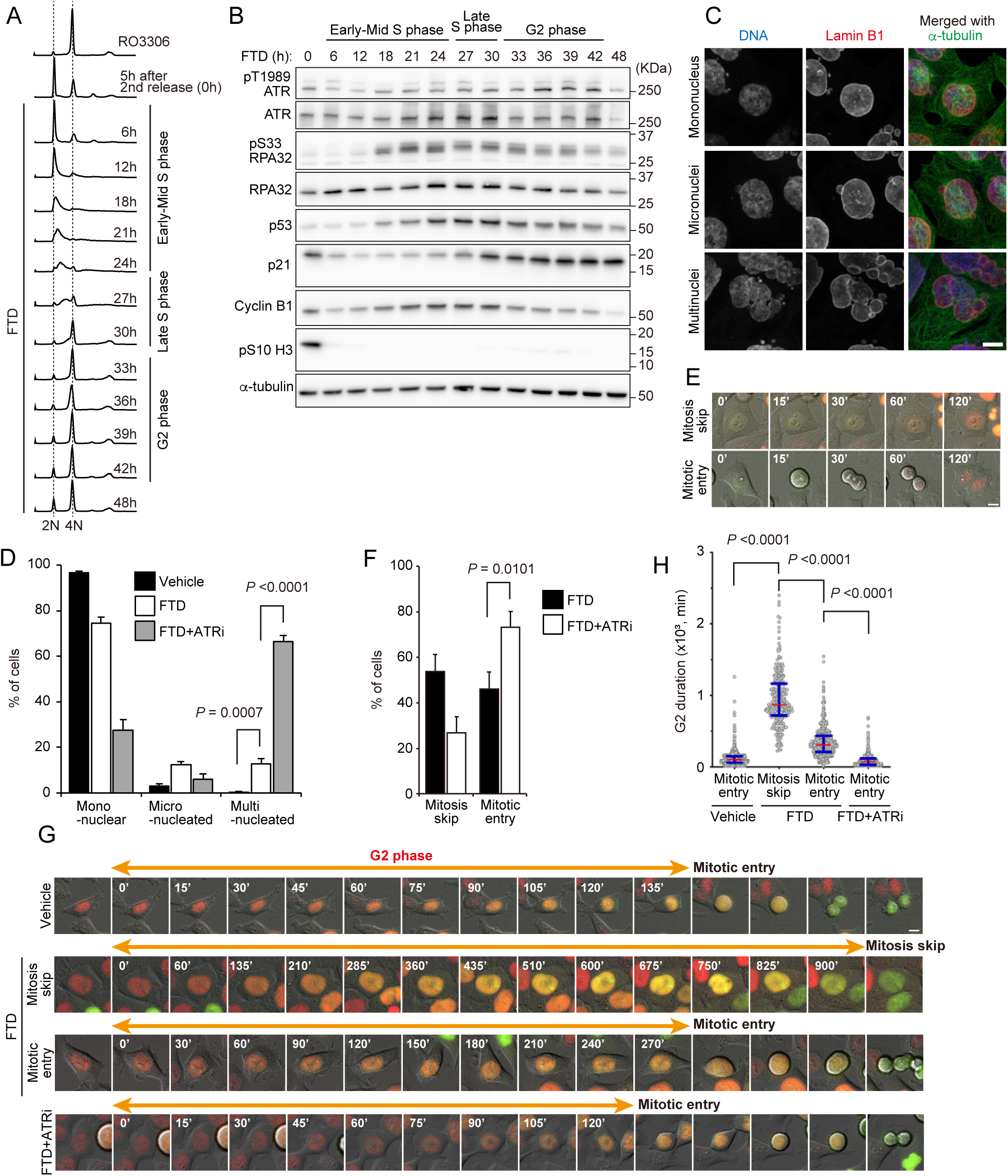
Activation of ATR and prolongation of G2 phase during TIS in p53-wild-type cells. **A**, **B.** Cell cycle analysis by flow cytometry and western blotting. Synchronous HCT116 cells were treated with FTD and then collected at indicated times after the second release. **A.** Propidium iodide (PI) staining was used to assess DNA content. **B.** Western blotting using antibodies specific for the indicated proteins. **C**. HCT116 cells expressing PIP-Fucci were treated with FTD, followed by sorting based on Cdt1 (1–17)-mVenus/ mCherry-Geminin (1–110)-double-positive signals. Western blot analysis was carried out using antibodies specific for the indicated proteins. **D, E.** HCT116 cells were treated with FTD/ATRi, or with FTD alone, and then fixed and stained with the indicated antibodies. Representative images of the cells are shown in D. Quantification of cells with aberrant nuclei formation is shown in E. Data are expressed as the mean ± the s.d. from three independent experiments (≥ 600 cells per experiment). **F, G.** HCT116 cells expressing Fucci-SA were treated with FTD/ATRi, or with FTD alone. Selected live cell images of representative phenotypes are shown in F. The proportion of cells exhibiting mitosis skip and entry is shown in G. Data are expressed as the mean ± s.d. from three independent experiments (≥ 50 cells per experiment). **H, I.** Selected frames from live-cell imaging of representative HCT116 cells expressing PIP-Fucci, treated with FTD alone or with FTD/ATRi. The orange left-right arrow denotes cells in the G2 phase (in H). Quantification of the duration of the G2 phase is shown in I. Open circles represent individual cells (n ≥ 250) from three independent experiments. Red lines, median; blue bars, interquartile range. Scale bar, 10 µm.

### An ATRi acts synergistically with the DRS-inducing drug FTD to induce cytotoxicity

Next, we investigated the behavior of cells after fate switching from TIS to mitotic entry induced by ATR inhibition. Our previous studies demonstrate that FTD-treated p53-deficient cells enter mitosis but not TIS, and ultimately undergo apoptosis ^9,10^. Therefore, we tested the hypothesis that ATRi-mediated cell fate switching of FTD-treated cells from senescence to mitotic entry may contribute to cytotoxicity. As expected, the combination of FTD and ATRi led to increased phosphorylation of histone H3 serine 10 (a mitotic marker), γH2AX (a DSB marker), and cleaved PARP/caspase3 (an apoptotic marker); this did not occur in cells treated with FTD alone (Fig. 2A and supplementary Fig.3A). Furthermore, combination treatment with the ATRi resulted in synergistic inhibition of FTD-treated cell growth (Fig. 2B). To confirm that this synergistic effect was due to cell death, we assessed the time course of cell death using dead cell detection methods. Consistent with our observation of apoptosis (Fig. 2A), accumulation of dead cells was much faster in the presence of FTD/ATRi than in that of FTD alone (Fig. 2C). Taken together, these results demonstrate that ATRi-induced cell fate switching from TIS to mitotic entry acts synergistically with FTD to induce cytotoxicity in p53 wild-type cells, which then undergo cell death in a manner similar to that of FTD-treated p53-deficient cells.

**Figure 2.**
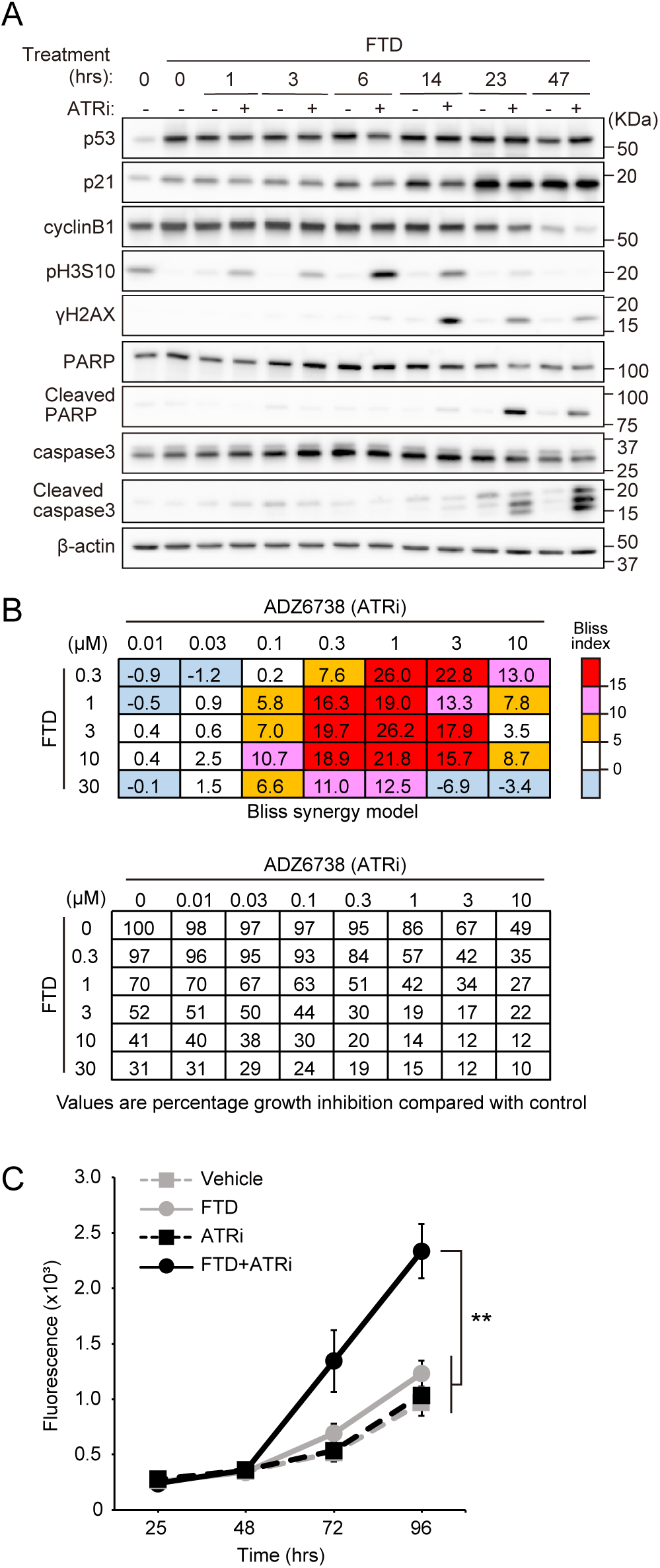
Synergistic cytotoxicity exerted by an ATR/FTD in p53-wild-type cells. **A.** HCT116 cells were treated with FTD for 25 h, followed by addition of the ATRi for an indicated times. Western blot analysis was carried out using antibodies specific for the indicated proteins. **B.** HCT116 cells ATRi were treated for 72 h with FTD in a 6×8 concentration grid. The Bliss additivism model was then used to classify additive and synergistic effects (see the Materials and methods section for details). **C.** Cell viability was analyzed using the CellTox Green Cytotoxicity Assay, which measures changes in membrane integrity that occur as a result of cell death in the presence of the indicated drugs. Data are expressed as the mean ± s.d. from three independent experiments.

### Temporal control of G2 phase by G2/M checkpoint inhibitors alters cell fate from TIS to mitotic entry

ATR plays crucial and multifaceted roles in the DRS response; indeed, one canonical function of the ATR pathway is that of a cell cycle checkpoint factor that arrests cells at G2 phase ^3,14^. Thus, we investigated whether the cell fate switch from TIS to mitotic entry induced by ATR inhibition is attributable to suppression of the G2/M checkpoint, which is positively regulated by ATR, CHK1, and WEE1 kinases^3^. To this end, we performed live-cell imaging to observe behavior of FTD-treated HCT116 Fucci-SA or PIP-Fucci cells in response to CHK1 and WEE1 kinase inhibitors, and found that these inhibitors phenocopied the effects of ATRi (e.g., cell fate switching from TIS to mitotic entry; Fig. 3A, and shortening of the G2 phase; Fig. 3B). Furthermore, we predicted that the shortening G2 duration in FTD-treated cells caused by ATR inhibition might be reversed by a PLK1 kinase inhibitor because activation of PLK1 is necessary for G2-M transition^15^. Indeed, additional inhibition of PLK1 slightly, but significantly, prolonged the duration of G2 (Fig. 3B). Similar to the effect of the ATRi, we found that a combination of FTD and a CHK1 or WEE1 inhibitor increased the number of DSBs (γH2AX) and the number of cells undergoing apoptosis (cleaved PARP/caspase3); this was not observed in cells treated with FTD alone (Fig. 3C, D). Consistent with these results, the dead cell detection assay revealed that accumulation of dead cells was more marked and faster after combined treatment with FTD and a CHK1 or WEE1 inhibitor than after treatment with FTD only (Fig. 3E, F).

**Figure 3.**
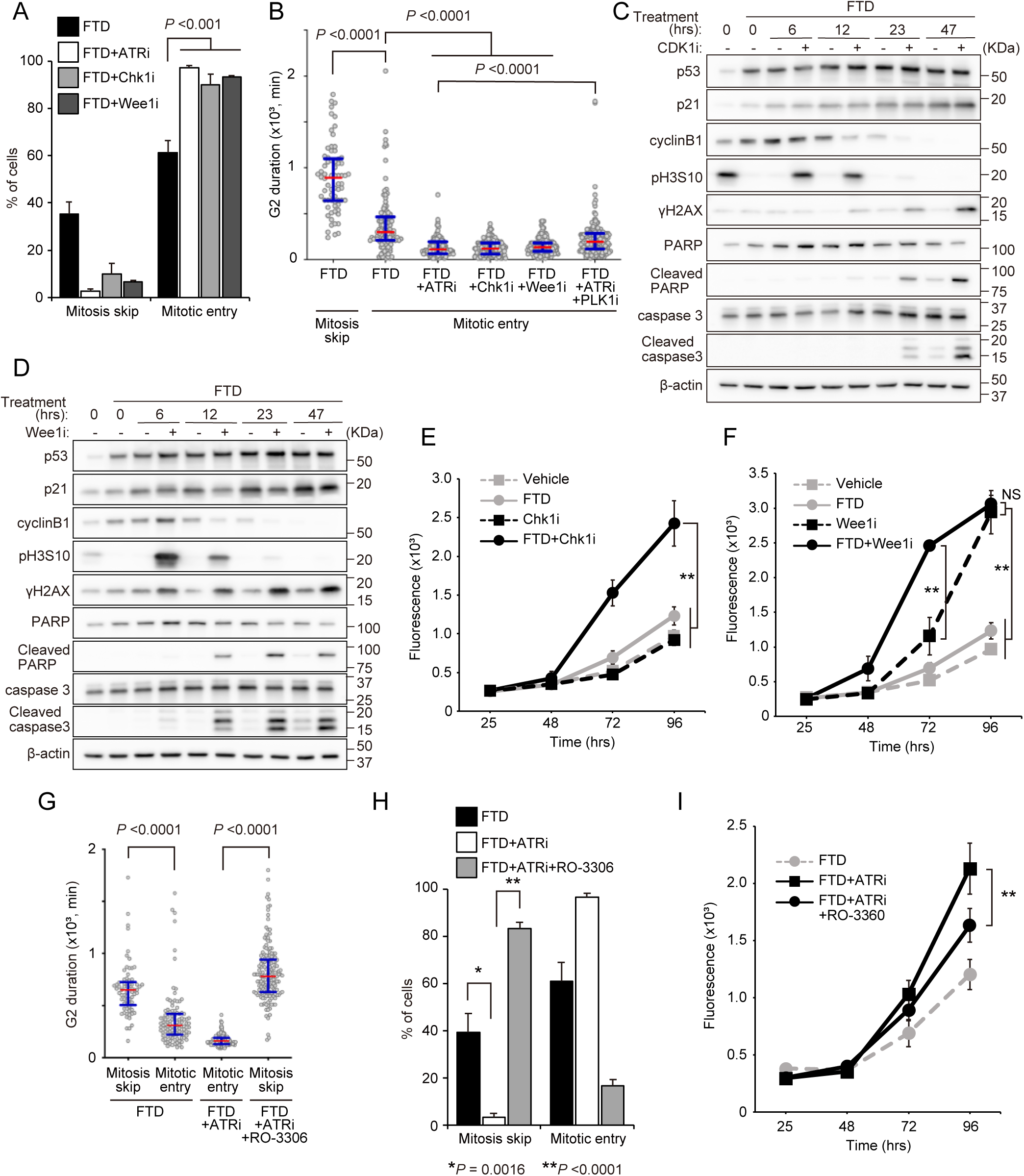
G2/M checkpoint inhibitors induce efficient cell death in FTD-treated p53-wild-type cells by shortening the G2 phase and allowing mitotic entry. **A.** HCT116 cells expressing Fucci-SA were treated with FTD plus G2/M checkpoint inhibitors, or with FTD alone and the proportion of cells exhibiting mitosis skip and entry was calculated. Data are expressed as the mean ± s.d. of three independent experiments (≥ 55 cells per experiment). **B.** The duration of the G2 phase was measured by live imaging of HCT116 cells expressing PIP-Fucci (n ≥ 55) and treated with FTD alone or with FTD/G2/M checkpoint inhibitors from three independent experiments. Open circles represent individual cells. Red lines, median; blue bars, interquartile range. **C, D.** HCT116 cells were treated with FTD for 25 h, followed by CHK1i (C) or WEE1i (D) for the indicated times. Western blot analysis was carried out using antibodies specific for the indicated proteins. **E, F.** Cell viability was analyzed using the CellTox Green Cytotoxicity Assay after treatment with the indicated drugs. Data are expressed as the mean ± s.d. from three independent experiments. **G.** Quantification G2 phase duration using live-cell imaging of HCT116 cells expressing PIP-Fucci (n ≥ 70) and treated with FTD alone, FTD/ATRi, or FTD/ATRi/CDK1i (RO-3306) from three independent experiments. Open circles represent individual cells. Red lines, median; blue bars, interquartile range. **H.** HCT116 cells expressing Fucci-SA were treated with FTD alone, FTD/ATRi, or FTD/ATRi/CDK1i. The proportion of cells exhibiting mitosis skip and entry is shown. Data are expressed as the mean ± s.d. from three independent experiments (≥ 170 cells per experiment). **I.** Cell viability was analyzed using the CellTox Green Cytotoxicity Assay after treatment with the indicated drugs. Data are expressed as the mean ± s.d. from three independent experiments.

Next, we investigated whether the duration of the G2 phase *per se* is important for determining cell fate switching of FTD-treated cells. As described above, DNA damage-induced mitosis skip is mediated by activation of p53-p21 activation, followed by premature activation of APC/C^Cdh1^, during G2 phase, resulting in degradation of Cyclin B1 and mitotic regulators ^7,8^. These data led us to hypothesize that a sufficient duration of G2 phase is required for mitosis skip via degradation of Cyclin B1 and mitotic regulators. Therefore, we asked whether artificial prolongation of the shortened G2 duration by the ATRi could restore mitosis skip. To answer this question, we measured G2 duration in the FTD/ATRi-treated PIP-Fucci cells following treatment (or not) with RO-3306, a CDK1 inhibitor that prevents G2-M transition, and confirmed that RO-3306 prolonged the shortened G2 duration (Fig. 3G). Accordingly, addition of RO-3306 altered the fate of FTD/ATRi-treated cells from mitotic entry to mitosis skip (Fig. 3H). Moreover, accumulation of dead FTD/ATRi-treated cells fell significantly after addition of RO-3306 (Fig. 3I). We conclude that both activation of the p53-p21 pathway and G2/M checkpoint-dependent prolongation of G2 duration by ATR are indispensable for FTD-induced senescence via mitosis skip, and thus that inhibition of G2/M checkpoint factors can trigger cell fate switching, resulting in cytotoxicity.

### Synergistic cytotoxicity of G2/M checkpoint inhibitors and FTD is independent of p53 status

Our recent study revealed activation of homologous recombination (HR) repair pathways in FTD-treated p53-KO cells during G2 phase, as evidence by formation of γH2AX and RAD51 foci^10^. Hence, the duration of G2 phase, controlled by the ATRi, is expected to affect the behavior not only of FTD-treated p53 wild-type cells, but also that of p53-KO cells. Thus, we examined whether the shortening of G2 duration in HCT116 p53-KO cells using G2/M checkpoint inhibitors acts synergistically to inhibit growth after treatment with FTD. Similar to the results observed for wild-type p53HCT116 cells, cells treated with FTD/ATRi exhibited higher levels of DSBs (γH2AX) and apoptosis (cleaved PARP/caspase3) than FTD-treated cells (Fig. 4A), as well as synergistic growth inhibition (Fig. 4B). Live-cell imaging of p53-KO cells indicated that the G2 phase was significantly longer in FTD-treated cells than in non-treated or FTD/ATRi-treated cells (Fig. 4C). Importantly, although the FTD/ATRi-treated p53 wild-type cells underwent cell death via a mechanism involving a cell fate switching from senescence to mitotic entry, followed by apoptosis, p53-KO cells also exhibited increased cytotoxicity even though the cell fate switch did not occur (Fig. 4D). Additional treatment with RO-3306 suppressed cytotoxicity induced by FTD/ATRi to levels similar to those seen in RO-3306-treated p53 wild-type cells induced to undergo artificial prolongation of the shorted G2 phase (Fig. 3G–I, 4E).

**Figure 4.**
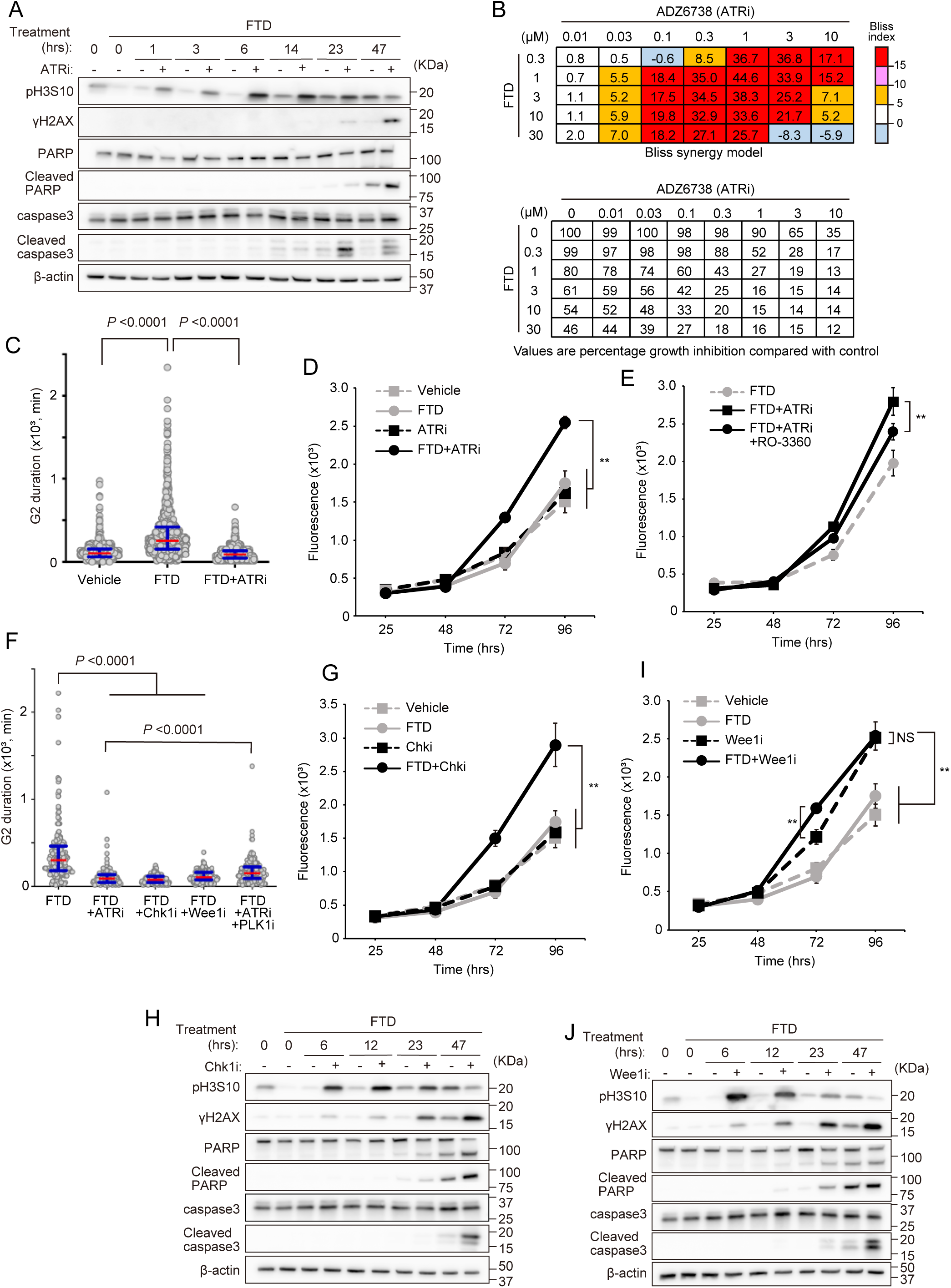
Shortening of G2 and G2/M checkpoint inhibitors act synergistically to induce cytotoxicity in FTD-treated p53-KO cells. **A.** HCT116 p53 KO cells were treated with FTD for 25 h, followed by the ATRi for the indicated times. Western blot analysis was carried out using antibodies specific for the indicated proteins. **B.** HCT116 p53 KO cells were treated with FTD/ATRi in a 6×8 concentration grid for 72 h. The Bliss additivism model was used to classify additive and synergistic effects (see Materials and methods section for details). **C.** Quantification of G2 phase duration using live-cell imaging ofHCT116 p53 KO cells expressing PIP-Fucci (n ≥ 900) and treated with FTD alone or FTD/ATRi from three independent experiments. Open circles represent individual cells. Red lines, median; blue bars, interquartile range. **D, E.** Cell viability was analyzed using the CellTox Green Cytotoxicity Assay after treatment with the indicated drugs. Data are expressed as the mean ± s.d. from three independent experiments. **F.** Duration of the G2 phase was measured using live-cell imaging ofHCT116 p53 KO cells expressing PIP-Fucci (n ≥ 160) and treated with FTD alone or FTD/G2/M checkpoint inhibitors from three independent experiments. Open circles represent individual cells. Red lines, median; blue bars, interquartile range. **G, I.** Cell viability was analyzed using the CellTox Green Cytotoxicity Assay after treatment with the indicated drugs. Data are expressed as the mean ± s.d. from three independent experiments. **H, J.** HCT116 p53 KO cells were treated with FTD for 25 h, followed by CHKi or WEE1i for the indicated times. Western blot analysis was carried out using antibodies specific for the indicated proteins.

Next, we confirmed that the phenomena induced in p53-KO cells by ATRi treatment could be replicated by a CHK1 or WEE1 inhibitor. As expected, cells treated with a CHK1 or WEE1 inhibitor also exhibited a shortening of the FTD-induced prolonged G2 phase. Furthermore, additional inhibition of PLK1 prolonged the G2 duration induced by FTD/ATRi (Fig. 4F). Consistent with these results, FTD-treated cells exhibited increased DSBs (γH2AX), apoptosis (cleaved PARP/caspase3) and cell death after additional treatment with a CHK1 (Fig. 4G, H) or WEE1 inhibitor (Fig. 4I, J). To clarify the generality of synergistic growth inhibition in FTD/ATRi-treated cells, we repeated the same experiments using cell lines other than HCT116. Both FTD-treated RKO (p53 wild type; Supplementary Fig 2A–C) and SW620 (p53 R273H; Supplementary Fig 2D–F) cell lines exhibited synergistic growth inhibition, apoptosis, and subsequent cell death after additional treatment of ATRi, suggesting that ATRi-induced cytotoxicity independent of genetic background.

### Combination of various DRS-inducing drugs and an ATRi increases cytotoxicity by shortening G2 phase

The data presented above show that tumor cells are sensitized to FTD-induced apoptotic cell death by addition of an ATRi, and that the mechanism involves shortening of the G2 phase. Furthermore, the effects of DRS-inducing drugs such as topoisomerase inhibitors, DNA-crosslinkers, and nucleoside analogues are potentiated by combination with an ATRi ^16–20^. To ascertain whether combined treatment with an ATRi and DRS-inducing drugs other than FTD shortens G2 and subsequently increases the number of dead cells, we performed live-cell imaging to measure the duration of the G2 phase in HCT116 p53-KO PIP-Fucci cells. In the previous experiments, the ATRi was added 25 h after addition of FTD to restrict ATR-mediated inhibition to the G2 phase; however, in these new experiments, the ATRi and DRS-inducing drugs, including FTD, were added to cells at the same time (this is because when numerous cells were just before entering G2 phase was different among DRS-inducing drugs). First, to determine the effect of the ATRi during G2 phase, we evaluated whether treatment of cells with DRS-inducing drugs at the IC_50_ concentration enter G2 phase without being continuously arrested in S phase. To this end, we treated cells with FTD, camptothecin (CPT), etoposide (ETO), doxorubicin (DOX), gemcitabine (GEM), or cisplatin (CDDP) and then measured the percentage of cells that entered G2 phase between 12 and 60 h post-treatment. We found that >90% of cells treated with FTD, CPT, ETO, or DOX treatment entered G2 phase, whereas the majority of cells treated with GEM or CDDP remained in S phase (Fig. 5A). Therefore, we then compared the G2 duration of cells treated with FTD, CPT, ETO, and DOX alone with that of cells treated with these compounds plus the ATRi. Similar to the results observed for FTD-treated cells, combination of drugs (CPT, ETO, and DOX) with the ATRi shortened G2 duration significantly; indeed, G2 duration was the same as that in unchallenged cells (Fig. 5B). Accumulation of dead cells increased markedly and more quickly after combined treatment than after monotherapy (Fig. 5C). Although the majority of GEM and CDDP-treated cells remained in S phase, the combination of ATRi plus GEM or CDDP triggered cell death, suggesting the ATRi affects cells in phases other than G2.

**Figure 5.**
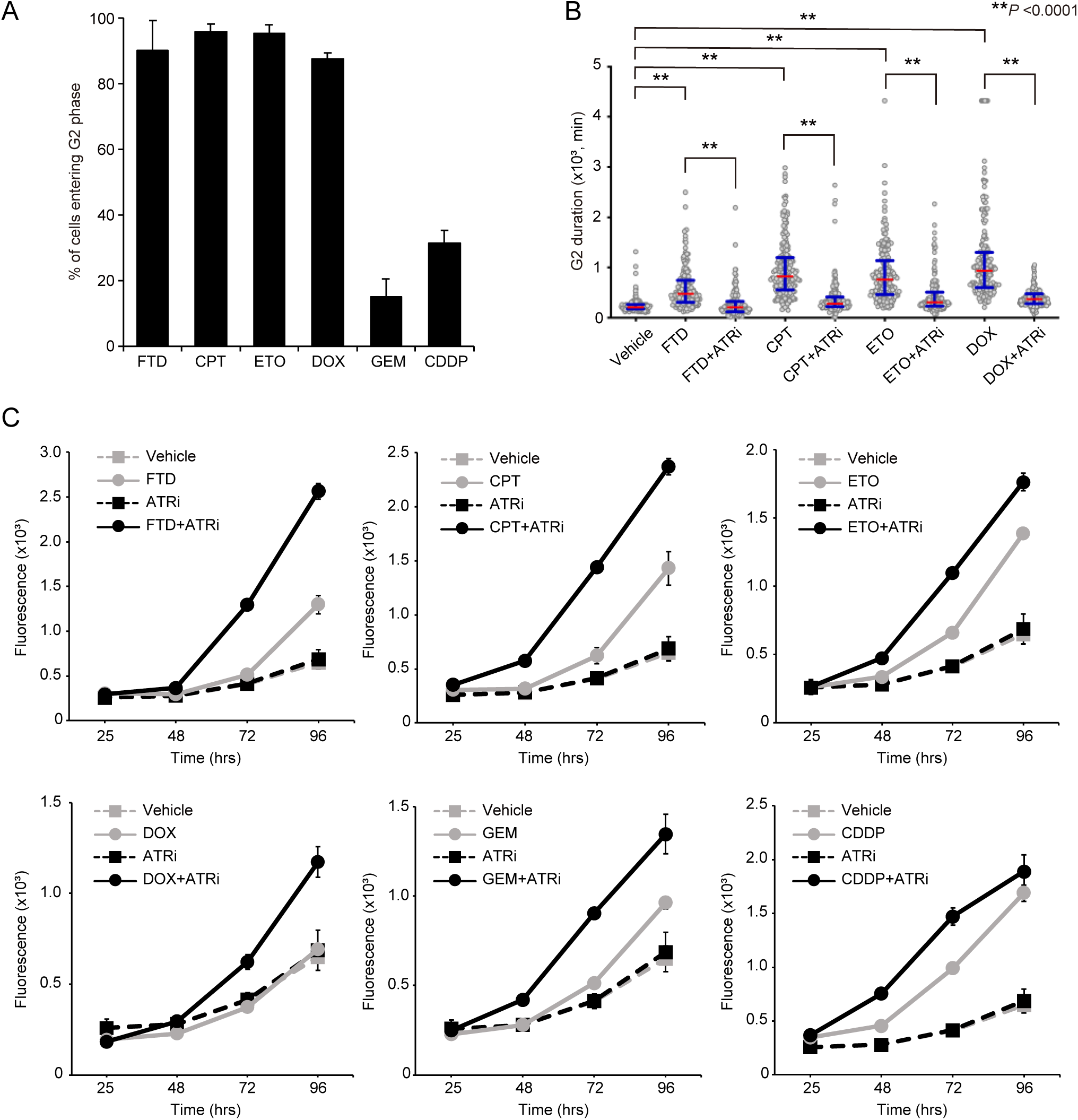
Effect of ATR inhibitors plus various DRS-inducing drugs on p53-KO cells. **A.** HCT116 p53 KO cells expressing PIP-Fucci were treated with FTD, camptothecin (CPT), etoposide (ETO), doxorubicin (DOX), gemcitabine (GEM), or cisplatin (CDDP), and the percentage of cells that entered G2 phase between 12 and 60 h after drug treatment was measured. Data are expressed as the mean ± s.d. from three independent experiments (≥ 60 cells per experiment). **B.** Duration of the G2 phase was measured using live-cell imaging of HCT116 p53 KO cells expressing PIP-Fucci (n ≥ 170) and treated with the indicated drugs from three independent experiments. Open circles represent individual cells. Red lines, median; blue bars, interquartile range. **C.** Cell viability was analyzed using the CellTox Green Cytotoxicity Assay after treatment with the indicated drugs. Data are expressed as the mean ± s.d. from three independent experiments.

Collectively, these results indicate that the enhanced cytotoxicity induced by FTD/ATRi, as well as other DRS-inducing drugs, may be caused by a shortening of G2 duration.

### ATRi contributes to increased formation of abnormal chromosome structures and DNA DSBs in FTD-treated cells during anaphase

Our recent report shows that FTD causes chromosomal abnormalities such chromosomal elongation and segmentation, possibly due to homologous recombination (HR) intermediates^21,22^ (Fig. 6A), and subsequent formation of numerous chromosome bridges during mitosis^10^. Therefore, we assessed the effects of ATR inhibition on FTD-induced chromosomal abnormalities. To this end, we analyzed chromosome spread in cells treated with FTD/ATRi or with FTD alone, and found that the ATRi increased the percentage of cells with more than 10 segmented chromosomes per cell significantly (Fig. 6A, B). To quantify the chromosome segmentation phenotype in detail, we performed fluorescence *in situ* hybridization (FISH) to label the centromeres of chromosome 3, 7 and 17 with probes, followed by DAPI staining. We then counted the number of segments in each labeled chromosome (Fig. 6C). Using this scoring method, we found FTD/ATRi increased the number of segments to one or more (S≥1) on chromosomes 7 and 17, and to two or more (S≥2) on chromosomes 3 and 7, compared with FTD alone (Fig. 6D). Furthermore, combination treatment increased the number of chromosome segments on chromosomes 3 and 7 significantly (Fig. 6E). In a previous study, we reported that FTD-induced chromosome segmentations result in chromosome bridges, including DNA bridges (which can be visualized by DAPI staining) and RPA-decorated ultra-fine DNA bridges (UFBs), which may be derived from HR intermediates^21^, as well as subsequent accumulation of DNA damage at the next G1 phase ^10^. Therefore, we speculated that synergistic cytotoxicity in FTD/ATRi-treated cells may occur via mitotic progression. To investigate this, we prepared early- or mid-mitotic cells by mitotic shake-off after treatment with nocodazole (Supplementary Fig.3D). In these cells, FTD/ATRi did not increase the number of DSBs (γH2AX), suggesting ATRi-induced synergistic effects do not occur during prometaphase (Fig. 6F). Next, to test another hypothesis (i.e., that the synergistic effect occurs during late mitosis), we obtained G1 cells that artificially escaped from chromosome segregation during anaphase to telophase after combined treatment with FTD/ATRi. Forced escape of FTD/ATRi-treated cells from late-mitosis was triggered using an aurora B kinase inhibitor (hesperadin) that induces transition from prometaphase to G1 phase without late-mitosis, a phenomenon called mitotic slippage ^23,24^. To more rigorously collect cells in G1 phase, we further sorted mitotic slipped-cells using Fucci-SA red (Supplementary Fig. 3E), and found that the number of DSBs (γH2AX) induced by FTD/ATRi was clearly lower in mitotic slipped-G1 cells than in G1 cells that passed through late mitosis, strongly suggesting that the ATRi-induced synergistic effect occurs during anaphase (Fig. 6G). We note that the difference in γH2AX levels between mitotic slipped-G1 cells and post-late mitotic G1 cells does not result from induction of apoptosis because post-mitotic G1 cells were collected early after the G1 phase transition, when no cleaved PARP was observed. Consistent with our previous study^10^, we found that FTD-treated cells exhibited abnormal nuclear morphologies, such as micronuclei and multinuclei. Furthermore, FTD/ATRi led to a significant increase in the number of multinuclear cells (Fig. 6H). As indicated above, FTD/ATRi-treated cells exhibited an increased number of DSBs (γH2AX) and higher levels of apoptosis than FTD-treated cells (Fig. 2A and 4A). Hence, we examined γH2AX foci in FTD/ATRi-treated cells, and found that FTD/ATRi increased formation of pan-nuclear γH2AX markedly, whereas formation of primary nuclear γH2AX foci decreased (Fig. 6I, J). Taken together, these data indicate that ATRi/FTD efficiently induces abnormal elongated- and segmented-chromosomes, resulting in more DNA damage via anaphase progression and, ultimately, efficient apoptosis.

**Figure 6.**
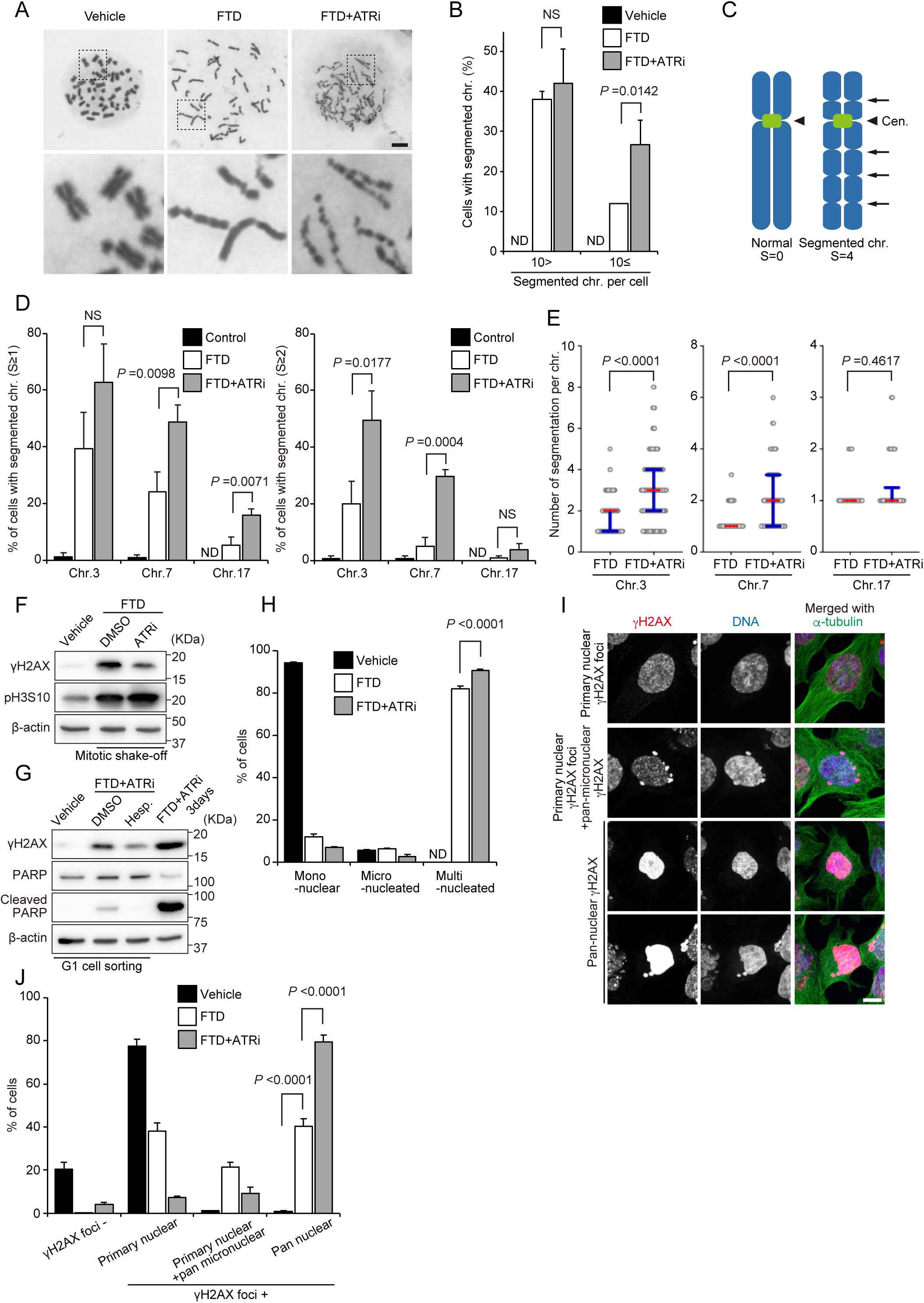
An ATR inhibitor increases formation of FTD-induced aberrant chromosomes, as well as DNA damage, in p53-KO cells. A, **B.** Giemsa-stained chromosome spread in FTD or FTD/ATRi-treated cells. Representative images of normal or segmented chromosome phenotypes are shown in A. The upper panels show whole chromosome spreads, whereas the lower panels show the indicated sections at higher magnification. Quantification of cells with segmented chromosomes is shown in B. Data are expressed as the mean ± s.d. from three independent experiments (≥50 cells per experiment). **C–E.** FISH-stained chromosome spreads of chromosomes 3, 7, and 17 from FTD- or FTD/ ATRi-treated cells labelled with probes that detect centromeric DNA. A schematic illustration of an abnormal metaphase chromosome showing four indentations (S=4) is shown in C. D shows the percentage of cells with chromosomes showing more than one indentation (S≥1; left), or more than two indentations (S≥2; right). The number of indentations in the indicated chromosomes is shown in E. Open circles represent an individual chromosome (≥150; chromosome 3, ≥90; chromosome 7, ≥20; chromosome 17) from three independent experiments. Red lines, medians; blue bars, interquartile range. **F.** Western blot analysis of γH2AX levels in extracts from asynchronous or mitotic (shake-off) cells treated with FTD/ATRi or with FTD alone. **G.** Western blot analysis of γH2AX and cleaved PARP levels in extracts from G1-phase-sorted vehicle-, FTD/ ATRi-, or FTD/ ATRi and hesperadin (Aurora B kinase inhibitor)-treated cells. An extract from cells treated with FTD/ATRi for 3 days is shown as a positive control. **H.** HCT116 cells were treated with FTD/ATRi or with FTD alone, fixed, and co-stained with DAPI and anti-LaminB/α-tubulin antibodies, and cells with aberrant nuclei formation were counted. Data are expressed as the mean ± s.d. from three independent experiments (≥ 900 cells per experiment). **I, J.** HCT116 p53-KO cells were treated with FTD/ATRi or with FTD alone, fixed, and stained with the indicated antibodies. Representative images of cells are shown in I. The percentage of cells exhibiting γH2AX accumulation in the primary nucleus or micronuclei is shown in J. Data are expressed as the mean ± s.d. from three independent experiments (≥ 600 cells per experiment).

## Discussion

DRS-inducing drugs activate the p53-p21 pathway and induce cellular senescence^6,25^ via mitosis skip^8,9^. During G2 phase, the ATR-CHK1-WEE1 axis acts as a DNA damage checkpoint factor^3^. Here, we found that DRS-inducing drugs lengthen the duration of the G2 phase significantly, and demonstrate that this delay is regulated by the ATR-CHK1-WEE1 axis. We also show that the length of the G2 phase decides the fate, i.e., mitosis skip or cellular senescence, of p53-wild-type cells. Inhibiting the ATR-CHK1-WEE1 axis shortens the G2 phase, thereby switching cell fate from senescence to mitotic entry and subsequent apoptosis (Fig. 7, upper). Moreover, when p53-KO cells treated with in DRS-inducing drugs, inhibition of the ATR-CHK1-WEE1 axis acted synergistically to increase cytotoxicity by significantly shortening G2 duration (Fig. 7, lower). Thus, we suggest the possibility that temporal regulation of the G2 phase is a novel approach to inducing synergy with DRS-inducing drugs, independent of p53.

**Figure 7.**
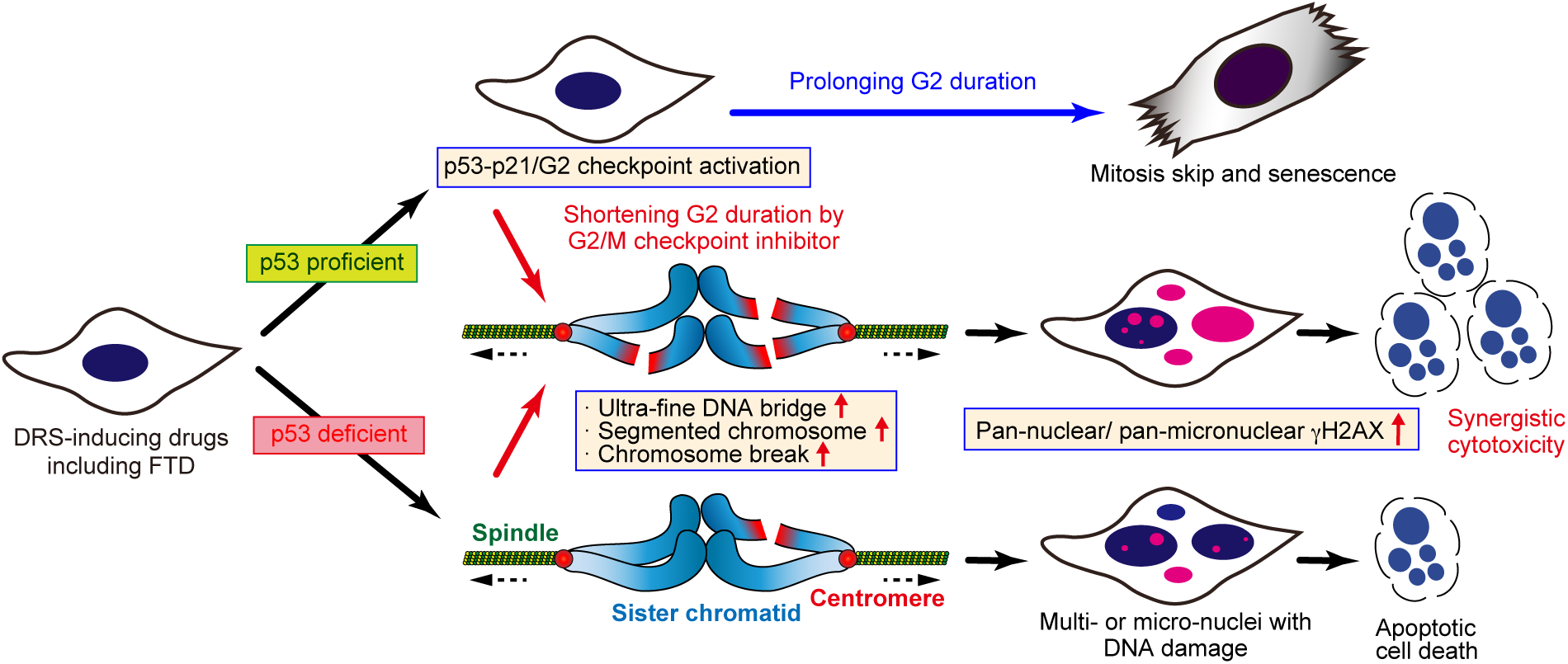
A model for mechanism of synergistic cytotoxicity of FTD and G2/M checkpoint inhibitors in a manner that is independent of p53 status.

Chemotherapeutic drugs, including DRS-inducing drugs, cause TIS^6^. Because TIS cancer cells arrest in G1 phase and stop proliferating, they can be considered a potential target for anticancer therapy. However, senescent cells still secrete inflammatory cytokines, chemokines, growth factors, and extracellular matrix remodeling factors that alter the local tissue environment and contribute to chronic inflammation and tumorigenesis. These phenomena come under the umbrella of a senescence-associated secretory phenotype (SASP)^26^. Our recent study showed that FTD induces secretion of SASP factors IL-1α, IL-1β, IL-8, TNFα, and CCL8 by HCT116 cells after transition into cellar senescence ^9,27^. Such SASP factors may promote tumor initiation and progression, along with chronic inflammation^28^. Furthermore, TIS cells can acquire cancer stemness and tumor aggressiveness when the activity of p53 is downregulated, resulting in subsequent escape from senescence, which can eventually result in cancer relapse and metastasis^29^. Therefore, our findings in this study, i.e., that prolonging G2 duration via activation of the ATR-CHK1-WEE1 axis is necessary for TIS transition, and that inhibition of the ATR-CHK1-WEE1 axis switches cell fate from TIS to cell death, avoid the above mentioned risks of TIS.

The mitosis skip seen in cells treated with IR or H_2_O_2_ and in cells expressing oncogenic RAS or undergoing replicative senescence ^7,8^ is also observed after FTD treatment. Although the shortening of G2 duration by ATRi switched cell fate from mitosis skip to mitotic entry, further inhibition of CDK1 kinase, which promotes G2-M transition, induced artificial arrest at G2 phase and restored prolonged G2 duration, mitosis skip, and subsequent cell proliferation (Fig. 3G–I). This suggests that DRS drugs-induced TIS requires concurrent degradation of Cyclin B1 and mitotic regulators through activation of the p53-p21 pathway, and temporal regulation of the G2 phase by activation of the ATR-CHK1-WEE1 axis. Although FTD treatment induced accumulation of ssDNA, and activation of ATR and p53-p21 during S phase, Cyclin B1 did not disappear immediately after entry into G2 phase (Fig. 1B), suggesting that a sufficient G2 duration is required for Cyclin B1 degradation and mitosis skip. Hence, cell fate (i.e., mitotic entry after DNA damage repair, or mitosis skip via activation of p53-p21 pathway-dependent degradation of Cyclin B1 and mitotic regulators) might be determined by the temporal threshold at G2. That is, if the mitosis skip threshold is breached first during a prolonged G2 phase, the cell skips mitosis and transits into cellar senescence; however, if DNA damage levels first fall below the G2/M checkpoint threshold as a result of repair, then cells enter mitosis. Our previous study of the mechanisms underlying FTD cytotoxicity demonstrate that FTD-induced DRS leads DNA damage, unresolved HR intermediates, chromosome bridges and, ultimately, apoptosis of p53-deficient cells ^9,10,30^. Because TIS cancer cells, which harbor wild-type p53, escape the cell cycle, arrest in G1, and become resistant to apoptotic stimuli via anti-apoptotic pathways, it is not hard to imagine that cell fate switching from mitosis skip to mitotic entry via inhibition of the ATR-CHK1-WEE1 axis may exert synergistic cytotoxic effects. However, the synergistic cytotoxic effects on FTD-treated cells exerted by inhibition of the ATR-CHK1-WEE1 axis were observed not only in p53-wild-type cells but also in p53-KO cells. We found that treatment with the ATRi increased structural chromosomal abnormalities, including elongated- and segmented-chromosomes, DSBs (γH2AX), and apoptosis in post-mitosis FTD-treated cells (Fig. 6). Since HR repair in FTD-treated cells occurs during G2 phase ^10^, the shortening of G2 duration induced by the ATRi might be associated with an increase in formation of segmented chromosomes. Whereas a prolonged G2 duration permits completion of HR repair at a certain frequency in the presence of FTD alone, shortening of G2 by combined treatment with FTD/ATRi does not allow enough time for HR repair, thereby increasing the number of chromosome abnormalities due to accumulation of HR intermediates. Combining various DRS-inducing drugs, including CPT, ETO and DOX, with the ATRi (Fig. 5C) led to increased cytotoxicity. Although the type of DNA damage responsible for increased cytotoxicity exerted by ATRi combination therapy differed according to the specific DRS-inducing drug, they may share a mechanism in which DNA damage is carried over into mitosis due to the shortened G2 phase. However, because the majority of cell treated with GEM or CDDP remained in S phase, it is possible that the increased cytotoxicity exerted by the ATRi/GEM or CDDP combinations could occur during intra-S phase, and via a mechanism different from that used by the FTD/ATRi combination.

In this study, the ATRi did not cause growth inhibition when used alone at the selected concentration (Fig. 2B, 4B; lower panels). Furthermore, a recent study reported that treatment with an ATRi alone does not affect proliferation of HT29 xenograft tumors ^31^. These findings suggest the possibility of an enhanced therapeutic effect with reduced off-target toxic side effects. Furthermore, our previous study concluded that FTD-induced genome instability rather than a catastrophic genome rearrangement, called chromothripsis (which is associated with aggressive tumor behavior; e.g., progression and chemoresistance, and a poor prognosis for cancer patients), was responsible for the increased cytotoxicity ^10^. Based on the results of this study, ATRi can be considered a modulator that increases the therapeutic efficacy of DRS-inducing drugs by promoting further genome instability. Because several small-molecule inhibitors of ATR, CHK1, and WEE1 kinases, are under clinical development^3,32^, further exploration of the mechanisms underlying synergistic effects of G2/M checkpoint inhibitors, including ATRi, and DRS-inducing drugs will be required before these agents can be tested in clinical practice.

### Materials and Methods Cell culture and reagents

HCT116 (ECACC), RKO (ATCC) and SW620 (ATCC) cells were cultured in DMEM (Thermo) All media were supplemented with 10% fetal bovine serum. All cell lines were grown in a 5% CO2 atmosphere at 37°C. All cell lines were authenticated by short tandem repeat (STR) analysis and were confirmed negative for *Mycoplasma* contamination with the MycoAlert Mycoplasma Detection Kit (Lonza). Fluorescent ubiquitination–based cell-cycle indicator (Fucci) SA or PCNA interacting protein (PIP) -Fucci cells were generated as described previously^10^. siRNA transfections were performed using Lipofectamine RNAiMax (Thermo) according to the manufacturer’s instructions. ATR-specific siRNA (5’- CCUCCGUGAUGUUGCUUGAdTdT-3’)^33^ and luciferase-specific siRNA (*GL3*; 5’- CUUACGCUGAGUACUUCGAdTdT -3’) were synthesized by Takara Bio and Sigma-Aldrich, respectively. A duplex targeting the luciferase gene (*GL3*) was used as a control. The final concentration of each reagent was as follows: trifluridine (FTD): 3 μM (T2511; Tokyo Chemical Industry), ceralasertib (AZD6738, ATRi): 200 nM (S7693; Selleck), rabusertib (LY2603618, CHK1i): 200 nM (S7693; Selleck), adavosertib (MK-1775, WEE1i): 200 nM (S1525; Selleck), BI 2536 (PLK1i): 5 nM (S1109; Selleck), RO-3306: 9 μM (SML0569; Sigma), camptothecin: 10 nM (208925; SIGMA), etoposide: 10 μM (E1383; SIGMA), doxorubicin: 300 nM (D1515; SIGMA), gemcitabine: 50 nM (G6423; SIGMA), cisplatin: 25 μM (D3371; Tokyo Chemical Industry), nocodazole: 200 ng ml^-1^ (M1404; SIGMA), hesperadin: 50nM (375680; Calbiochem).

### Western blotting

Cell pellets were sonicated with 1xSDS sample buffer [62.5 mmol/L Tris-HCl (pH 6.8), 2.5% SDS, 0.002% Bromophenol Blue, 5% β-mercaptoethanol, 10% glycerol and were boiled at 95°C for 5 min. Cell lysates were separated by SDS-PAGE with TGX precast gels (Bio-Rad) and transferred to membranes using the Trans-Blot Turbo Transfer System (Bio-Rad). Membranes were incubated with primary antibodies and horseradish peroxidase-conjugated secondary antibodies. Membranes were imaged using ImageQuant LAS 4000 mini (GE Healthcare). The antibodies used are listed in TableS1.

### Immunofluorescence

Cells were grown collagen-I-coated coverslips (IWAKI). Thereafter, cells were rinsed with PBS at 37℃, fixed in 4% paraformaldehyde for 15 min, permeabilized in PBS containing 0.1% Triton X-100 for 5 min, blocked in PBS containing 2% BSA for 30 min at room temperature, and incubated with the primary antibodies (Supplementary Table 3) and Alexa Fluor 488-, 568-, 594- and 647-conjugated secondary antibodies (Thermo) diluted 1:2000. Coverslips were washed with PBS containing 4,6-diamidino-2-phenylindole (DAPI) for 5 min, washed with PBS and mounted with Prolong Glass (Thermo).

### Image acquisition and analysis

For fixed-cell experiments, fluorescence image acquisition was performed using a Nikon A1R confocal imaging system controlled by the Nikon NIS Elements software (Nikon). The objective lens was an oil immersion Plan-Apo 100× numerical aperture (NA) 1.40 lens (Nikon). Images were acquired as Z-stacks at 0.2-µm intervals and deconvoluted. Maximum-intensity projections were generated using the NIS Elements software (Nikon). For live-cell imaging of cell-cycle progression, HCT116-Fucci, HCT116 p53^-/-^-Fucci, HCT116-PIP Fucci, HCT116 p53^-/-^-PIP Fucci cells were cultured in a glass bottom chamber (Matsunami) that had been coated with collagen-I (Nippi) containing FluoroBrite DMEM (Thermo) supplemented with 10% FBS and GlutaMAX (Thermo). Cells were maintained at 37 °C in the presence of 5% CO_2_ in a stage-top incubator and PureBox SHIRAITO (Tokai Hit). Images were acquired every 10 min using a Plan-Apo 20× NA 0.75 lens (Nikon) on an inverted fluorescence microscope Eclipse Ti-E (Nikon) equipped with a DS-Qi2 camera (Nikon). For chromosome spreading experiments, an oil immersion Plan-Apo 100× numerical aperture (NA) 1.40 lens (Nikon) on a BZ-X800 fluorescence microscope (KEYENCE) was used for image acquisition.

### Cell viability assay

Cell viability was determined using CellTiter Glo assay (Promega). Briefly, cells were seeded at 4,000 cells/well in a 96-well plate. After 24 hours, varying concentrations of drug were dosed to the cell, either as a single agent or in combination (FTD 0-100μM, ATRi 0-10μM). The luminescence signal for CellTiter Glo was determined 72 hours after dosing.

The Bliss additivism model was used to classify the additivity and synergy effect of combining FTD and ATRi as additive, synergistic, or antagonistic. A theoretical curve was calculated for combined inhibition using the equation Ebliss = EA + EB - EA x EB, where EA and EB are the fractional inhibitions obtained by drug A alone and drug B alone at specific concentrations. Here, Ebliss is the fractional inhibition that would be expected if the combination of the two drugs was exactly additive. If the experimentally measured fractional inhibition is less than Ebliss, the combination was said to be synergistic. If the experimentally measured fractional inhibition is greater than Ebliss, the combination was said to be antagonistic^16^.

### Cell cytotoxicity assay

Cell cytotoxicity was assessed using the CellTox Green Cytotoxicity Assay (Promega), according to the manufacturers’ instructions. Briefly, cells were seeded at 4,000 cells/well in a 96-well plate. 24 hours after plating, each agents were added. After another 25hours, CellTox Green reagents were added with a newly medium containing the reagents. Fluorescence signals were measured consecutively for three days. The graph shows the quantification of the fluorescence intensity.

### Chromosome spreads and fluorescence *in situ* hybridization (FISH) analysis

Chromosome spreading and FISH analyses were performed as previously described ^34^. Briefly, cells were treated with 100 ng/ml KaryoMAX Colcemid Solution (Thermo). Each sample was collected and hypotonically swollen in 75 mM KCl for 20 min at 37°C when the percentage of mitotic cells was equivalent (FTD: for 30hr, the combination of FTD and ATRi: for 6hr). Cells were treated with hypotonic solution (0.075 M KCl) for 12 min at 37°C and fixed in Carnoy’s fixative solution (75% methanol and 25% acetic acid) with two changes of the fixative. Cells were dropped onto cooled glass slides and dried at 65°C for 60 sec. After then, for chromosome morphology observations, chromosomes were stained in 5% Giemsa for 15 min, rinsed with PBS, air-dried for one day and mounted using NEW MX (Matsunami). For identification of chromosomes, after cells were dropped onto cooled glass slides and dried described above, the coverslips were washed in 2× SSC (1× SSC is 0.15 M NaCl and 0.015 M sodium citrate) for 2 min, and then dehydrated in an ethanol series (70%, 85%, and 100%; 2 min each following air drying). The DNA FISH probes (ZytoLight SPEC p16/CEN3/7/17 Quadruple Color Probes; Zytovision) were pre-warmed at 37°C, placed onto the coverslips, and denatured at 75°C for 2 min. The coverslips were then incubated overnight at 37°C, washed with 4× SSC containing 0.05% Tween 20 for 5 min, and then incubated in 0.25× SSC at 72°C for 2 min. After a wash in 4× SSC containing 0.05% Tween 20 for 30 sec, the coverslips were mounted in ProLong Glass containing DAPI (Thermo).

### Statistical analysis

Sample numbers, *n* numbers and *P* values are described in the figure legends. Data were analysed using unpaired two-tailed Student’s *t*-test in figure 1E, 1G, 2C, 3A, 3F-F, 3H-I, 4D-E, 4G-I, 5A, 5C, 6B, 6D, 6H, 6J and supplementary figure 1C; Mann-Whitney *U*-test in figure 1I, 3B, 3G, 4C, 4F, 5B and 6E. The normality of the data and variance between the groups were tested. Statistical calculations were carried out using the PRISM program (GraphPad Software).

## Acknowledgements

We thank Masako Kosugi, Atsuko Yamaguchi, and Itsuki Sakai for expert technical assistance, as well as the staff at The Research Support Center, Research Center for Human Disease Modelling, Graduate School of Medical Sciences, Kyushu University, for technical support. This work was supported by the Ministry of Education, Culture, Sports, Science and Technology of Japan (awarded to K. Nonaka, JSPS KAKENHI grant number 25K12043 and H. Kitao, JSPS KAKENHI grant number 23K24030).

## Author contributions

M.I. conceptualized the study and contributed to the study design. K.N carried out most of the experiments, analyzed the data. M.I., T.Y., Y.M. and H.K. supervised the project. M.I. and K.N wrote the manuscript. H.K. edited the manuscript. T.W., H.O., Y.K., K.A. and E.O. assisted with data analysis and interpretation. All authors discussed the results and reviewed and approved the manuscript.

