## Supplemental Figures for "Temporal regulation of G2 phase avoids therapy-induced senescence caused by DNA replication stress-inducing drugs and provides synergistic cytotoxicity"

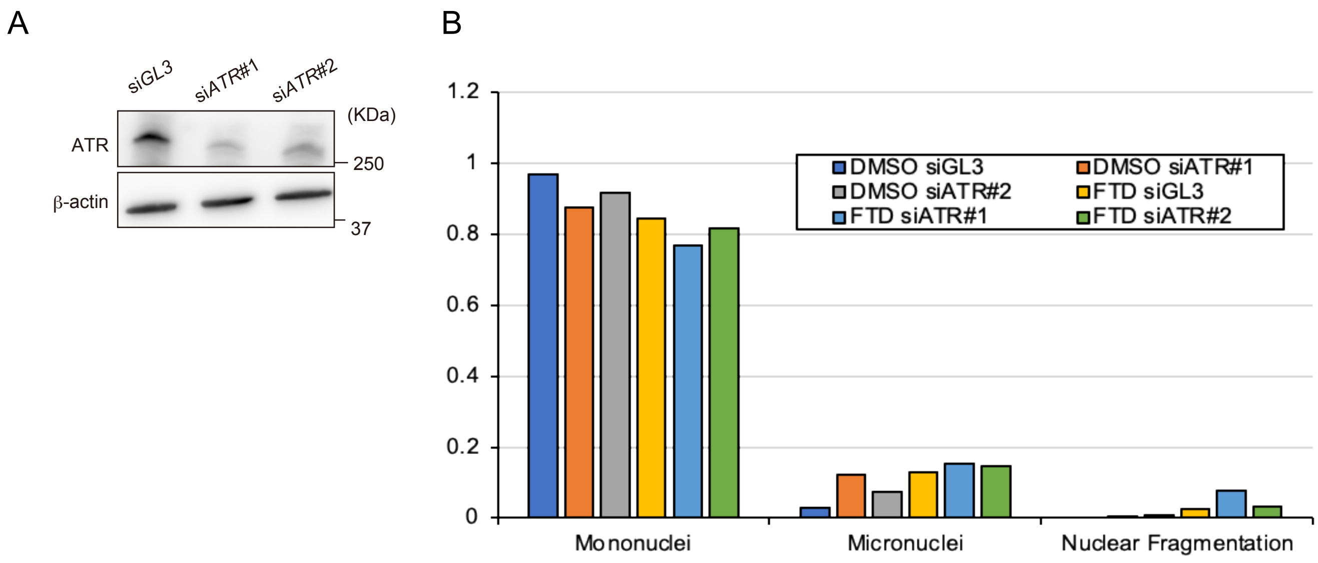
A
siGL3
siATR#1
siATR#2
(KDa)
ATR
250
β-actin
37
B
1.2
1
0.8
0.6
0.4
0.2
0
DMSO siGL3
DMSO siATR#1
DMSO siATR#2
FTD siGL3
FTD siATR#1
FTD siATR#2
Mononuclei
Micronuclei
Nuclear Fragmentation

Suppl. Fig. 1

A RKO (p53 Wild-type)

ADZ6738 (ATRi)

| FTD (μM) | 0.01 | 0.03 | 0.1 | 0.3 | 1 | 3 | 10 |
|---|---|---|---|---|---|---|---|
| 1 | -1.1 | -0.4 | 5.2 | 29.3 | 26.0 | -2.7 | -6.1 |
| 3 | 1.0 | 3.6 | 20.2 | 39.7 | 19.7 | -7.9 | -8.0 |
| 10 | 0.3 | 3.2 | 15.4 | 25.0 | 12.1 | -7.4 | -9.7 |
| 30 | 0.1 | 1.7 | 8.0 | 11.8 | 3.8 | -10.9 | -12.2 |
| 100 | 0.2 | 2.1 | 4.0 | 3.3 | -6.0 | -19.0 | -16.7 |

Bliss index: 15, 10, 5, 0

Bliss synergy model

ADZ6738 (ATRi)

| FTD (μM) | 0 | 0.01 | 0.03 | 0.1 | 0.3 | 1 | 3 | 10 |
|---|---|---|---|---|---|---|---|---|
| 0 | 100 | 99 | 99 | 99 | 96 | 70 | 24 | 13 |
| 1 | 84 | 84 | 84 | 78 | 51 | 33 | 23 | 17 |
| 3 | 66 | 64 | 62 | 45 | 23 | 26 | 24 | 16 |
| 10 | 47 | 46 | 43 | 31 | 20 | 21 | 19 | 16 |
| 30 | 30 | 30 | 28 | 22 | 17 | 17 | 18 | 16 |
| 100 | 20 | 19 | 17 | 15 | 15 | 20 | 23 | 20 |

Values are percentage growth inhibition compared with control

D SW620 (p53 R273H)

ADZ6738 (ATRi)

| FTD (μM) | 0.01 | 0.03 | 0.1 | 0.3 | 1 | 3 | 10 |
|---|---|---|---|---|---|---|---|
| 0.3 | 0.8 | 0.5 | -1.5 | 11.1 | 19.1 | 20.7 | 13.6 |
| 1 | 2.8 | 5.0 | 7.8 | 19.6 | 26.5 | 24.6 | 14.2 |
| 3 | 2.2 | 4.8 | 9.4 | 19.9 | 24.7 | 22.4 | 13,2 |
| 10 | 2.3 | 4.9 | 10.9 | 20.6 | 22.7 | 20.3 | 8.7 |
| 30 | 2.5 | 5.2 | 10.2 | 16.5 | 16.5 | -8.3 | -6.2 |

Bliss index: 15, 10, 5, 0

Bliss synergy model

ADZ6738 (ATRi)

| FTD (μM) | 0 | 0.01 | 0.03 | 0.1 | 0.3 | 1 | 3 | 10 |
|---|---|---|---|---|---|---|---|---|
| 0 | 100 | 101 | 101 | 99 | 94 | 79 | 69 | 51 |
| 1 | 63 | 63 | 63 | 64 | 48 | 30 | 22 | 18 |
| 3 | 56 | 54 | 52 | 49 | 33 | 18 | 14 | 14 |
| 10 | 48 | 47 | 44 | 39 | 25 | 13 | 11 | 11 |
| 30 | 44 | 43 | 40 | 33 | 21 | 12 | 10 | 10 |
| 100 | 41 | 39 | 36 | 30 | 22 | 16 | 13 | 11 |

Values are percentage growth inhibition compared with control

B

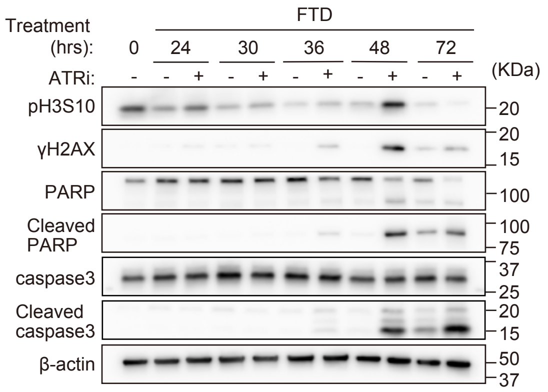


E

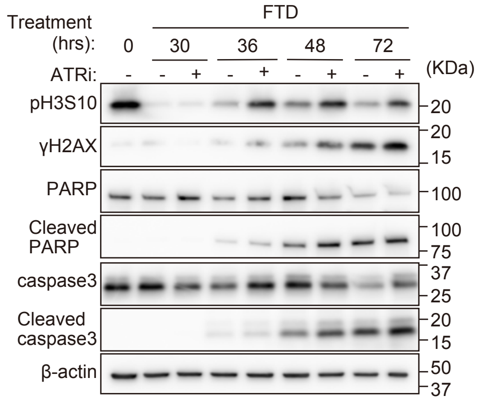


C

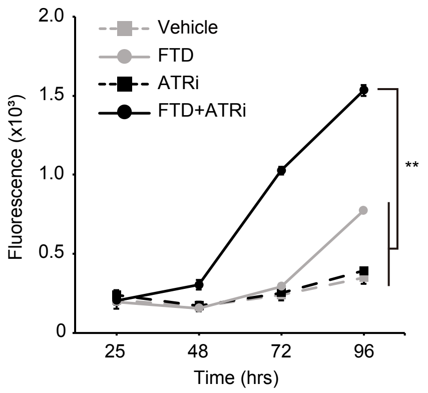


F

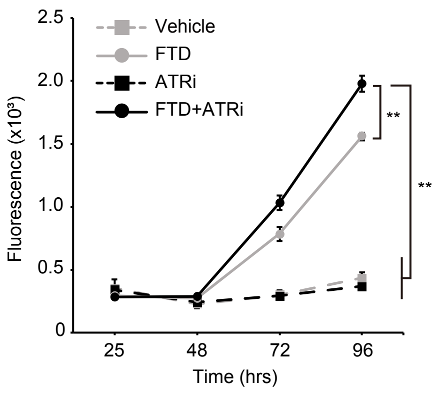


Suppl. Fig. 2

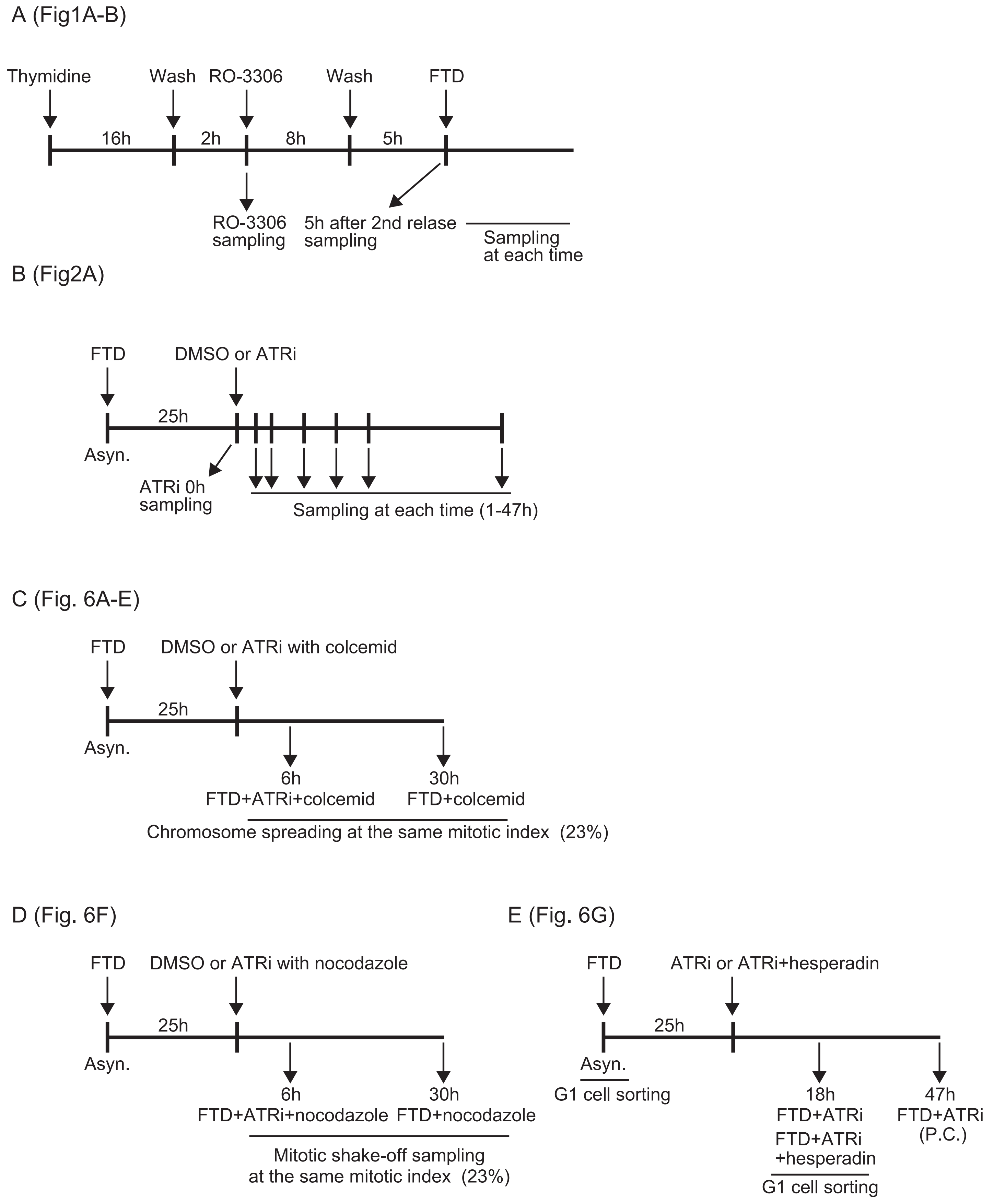

A (Fig1A-B)
Thymidine
Wash
RO-3306
Wash
FTD
16h
2h
8h
5h
RO-3306
sampling
5h after 2nd relase
sampling
Sampling
at each time
B (Fig2A)
FTD
DMSO or ATRi
25h
Asyn.
ATRi 0h
sampling
Sampling at each time (1-47h)
C (Fig. 6A-E)
FTD
DMSO or ATRi with colcemid
25h
Asyn.
6h
FTD+ATRi+colcemid
30h
FTD+colcemid
Chromosome spreading at the same mitotic index (23%)
D (Fig. 6F)
FTD
DMSO or ATRi with nocodazole
25h
Asyn.
6h
FTD+ATRi+nocodazole
30h
FTD+nocodazole
Mitotic shake-off sampling
at the same mitotic index (23%)
E (Fig. 6G)
FTD
ATRi or ATRi+hesperadin
25h
Asyn.
G1 cell sorting
18h
FTD+ATRi
FTD+ATRi
+hesperadin
G1 cell sorting
47h
FTD+ATRi
(P.C.)


Suppl. Fig. 3
